# Regulating Light-Harvesting Protein Assembly through Engineered Trimers of Phycocyanin and Allophycocyanin

**DOI:** 10.64898/2026.06.24.734401

**Authors:** Motoyasu Adachi, Masaaki Tsubouchi, Chie Shibazaki, Keita Miyake, Ryuji Itakura

**Affiliations:** Institute for Quantum Life Science, National Institutes for Quantum Science and Technology (QST), 4-9-1 Anagawa, Inage, Chiba 263-8555, Japan; Kansai Institute for Photon Science, National Institutes for Quantum Science and Technology (QST), 8-1-7 Umemidai, Kizugawa, Kyoto 619-0215, Japan; J-PARC Center, Japan Atomic Energy Agency (JAEA), 2-4 Shirakata, Tokai, Naka, Ibaraki 319-1195, Japan; Department of General Systems Studies, Graduate School of Arts and Sciences, The University of Tokyo, 3-8-1 Komaba, Meguro, Tokyo, 153-8902 Japan

**Keywords:** Phycobiliprotein, Phycocyanin, Allophycocyanin, Protein engineering, X-ray crystallography, Trimer, Light harvesting

## Abstract

Phycobilisome-associated phycobiliproteins naturally assemble into trimeric and hexameric architectures, making it difficult to isolate stable trimeric species for structural and functional studies. Here, we establish a rational strategy to generate stable trimers of phycocyanin (TeCPC) and allophycocyanin (TeAPC) from *Thermosynechococcus elongatus* by destabilizing the hexamerization interface. We introduce structure-guided glycine-to-arginine substitutions, G29R in TeCPC and G21R in TeAPC, to suppress hexamer formation. In TeCPC, an additional T121Y substitution in the α-subunit further stabilizes the trimeric assembly, which was identified through screening of structure-based variants. Recombinant expression in *Escherichia coli* produced holoproteins with native-like chromophorylation. Size-exclusion chromatography, mass spectrometry, and X-ray crystallography confirm the homogeneous trimers, with no evidence of hexamer formation under the conditions examined. Thermal analyses demonstrate melting temperature exceeding 69 °C for both engineered complexes. These results provide a general framework for obtaining stable phycobiliprotein trimers for structural and spectroscopic investigations, and for controlling phycobiliprotein quaternary structure.

## 1. Introduction

Phycobilisomes (PBS) are the major light-harvesting antenna complexes of cyanobacteria and red algae, which assemble from phycobiliproteins and linker proteins into massive functional units [1–3]. Within these assemblies, the structural hierarchy begins with α- and β-subunits that form αβ heterodimers. The fundamental quaternary structure is a trimeric ring composed of three αβ heterodimers. These trimers further associate face-to-face to form hexamers, representing a higher-order quaternary structure that constitutes the peripheral rods and the central core of the PBS architecture [4,5]. Because these architectures define the precise spatial arrangement of bilin chromophores within the protein scaffold, they are intrinsically linked to the highly evolved photophysical properties and excitation energy transfer dynamics of phycobiliproteins [6,7].

To elucidate the fundamental structure–function relationships and energy dynamics of these light-harvesting networks, highly uniform biophysical and structural models are essential. However, isolated phycobiliproteins frequently exhibit oligomeric plasticity, existing in an equilibrium between trimeric and hexameric states. This structural heterogeneity complicates the interpretation of mechanistic data, as variations in spectroscopic signals, thermal behavior, or crystal packing may reflect transient assembly states rather than intrinsic properties of the chromophore environment. Consequently, a system restricted to a well-defined quaternary structure—specifically the minimal functional trimer—is required to provide a reliable basis for high-resolution structural and spectroscopic studies.

Heterologous expression in *Escherichia coli* (*E. coli*) provides direct genetic access to the protein scaffolds, yet the faithful reconstruction of holo-phycobiliproteins requires the coordinated co-expression of biosynthetic enzymes to ensure native-like post-translational modifications [8]. Phycocyanobilin must be synthesized and covalently attached to specific cysteine residues by bilin lyases, and the conserved β-subunit γ-*N*-methylasparagine should also be installed by CpcM [9,10]. Achieving efficient chromophorylation while maintaining precise control over the quaternary structure remains a significant technical challenge in recombinant systems.

In this study, we employed a rational structural design approach to inhibit higher-order association and stabilize defined trimeric quaternary structures of phycocyanin (TeCPC) and allophycocyanin (TeAPC) from the thermophilic cyanobacterium *Thermosynechococcus elongatus* BP-1 [11]. Despite the importance of the trimer as the fundamental building block of phycobilisomes, stable isolated trimers are often difficult to obtain because of spontaneous hexamerization. We therefore sought to establish a simple and generally applicable strategy for preventing hexamer formation while preserving the structural integrity of phycobiliproteins. To obtain structurally homogeneous trimers, we introduced structure-guided glycine-to-arginine substitutions near the hexamer interface (G29R in TeCPC and G21R in TeAPC) to suppress higher-order assembly. For TeCPC, a trimer-promoting mutation (T121Y) was additionally identified through screening of structure-based variants and incorporated into the final construct. We characterized these engineered complexes using size-exclusion chromatography, X-ray crystallography, mass spectrometry, and thermal analysis to verify their oligomeric homogeneity and the fidelity of their post-translational modifications.

The resulting engineered TeCPC and TeAPC variants generated in this study provide structurally and biochemically defined model systems suitable for future investigations into phycobiliprotein function. By decoupling the functional trimer from higher-order assembly while maintaining native-like chromophore attachment, these complexes facilitate structural and spectroscopic studies [12].

## 2. Materials and Methods

### 2.1. Plasmid construction

All DNAs were chemically synthesized and purchased from Genewiz (Azenta Life Sciences, Japan). Their coding sequences were optimized for *E. coli* expression. Genes encoding TeCPC and TeAPC from *Thermosynechococcus elongatus* BP-1 were designed and inserted into the pET24a vector.

### 2.2. Site-directed mutagenesis

For the primary TeCPC (Fig. S1), a set of 28 primers and PrimeSTAR DNA polymerase were used for the sub-cloning by PCR method to prepare mutant genes (Table S1). The amplified PCR products were introduced into competent HST08 cells (Takara, Japan). Plasmids were prepared using QIAprep Spin Miniprep Kit (Qiagen). The DNA sequence coding for the proteins was confirmed by DNA sequencing on the Applied Biosystems SeqStudio Genetic Analyzer (Thermo Fisher Scientific).

### 2.3. Expression plasmids and chromophore incorporation

The recombinant proteins TeCPC G29R (additionally containing the T121Y substitution in the α-subunit) and TeAPC G21R were constructed using an *E. coli* expression system. Gly29 in TeCPC and Gly21 in TeAPC are located in the α-subunits.

Supplementary Table S2. lists the expression plasmids used in this study. The protein sequences involved in posttranslational modifications were obtained from previous reports [9,13–15]. The pACYC_HO1PcyA and pCDF_MTSEF were used for posttranslational modifications of primary TeCPC variants, TeCPC G29R, and TeAPC G21R. DNA and encoded amino acid sequences of proteins have been deposited into the DNA Data Bank of Japan. *E. coli* JM109(DE3) cells (Promega) were transformed with the expression plasmids to produce recombinant proteins.

### 2.4. Expression in E. coli for protein purification

The bacterial cells were grown in LB or Terrific Broth medium containing 15 mg/L kanamycin, 7 mg/L chloramphenicol, and 10 mg/L streptomycin. An overnight preculture using 20 mL LB medium at 310 K was inoculated into 400 mL of TB medium in a 3 L Erlenmeyer flask and incubated with shaking at approximately 90 rpm using a BR 1 incubator shaker (TAITEC). At an OD600 of approximately 0.7, protein expression was induced at 310 K for 72 hours by adding isopropyl-β-D-thiogalactopyranoside to a final concentration of 0.1 mM, as well as 5-aminolevulinic acid (final 80 mg/L), biotin (final 0.25 mg/L), vitamin B12 (final 0.135 mg/L), and thiamine (final 0.335 mg/L). In small-scale screening experiments for the primary TeCPC mutants, 3 mL of TB medium was used for protein expression.

### 2.5. Purification for TeCPC G29R and TeAPC G21R

The cells were harvested by centrifugation at 5000 g for 20 min. The harvested cells were suspended into Buffer T (20 mM Tris-HCl, pH 8.0) and disrupted by sonication. The crude extract was centrifuged at 10000 g for 30 min to collect the supernatant. The supernatant was dialyzed against 1 L of Buffer T overnight and then centrifuged at 10000 g for 30 min to remove the insoluble fraction. The obtained supernatant was loaded onto a column filled with Ni Sepharose 6 Fast Flow (Cytiva) at a bed volume of 20 mL and eluted with 0.5 M imidazole in Buffer T after washing with Buffer T. The eluted solution was applied onto a StrepTrap XT column with 5 mL volume (Cytiva) and eluted by 60 mg/mL biotin dissolved in 0.5 M Tris-HCl buffer (pH 8.0) with NaOH. The purified protein solution was subsequently dialyzed against Buffer T. The protein concentrations of TeCPC G29R and TeAPC G21R were estimated using UV absorption at 280 nm with molecular extinction coefficients of 108,990, and 101430 M^−1^ cm^−1^, respectively, as calculated from the amino acid content [16].

### 2.6. Size□exclusion column chromatography

The quaternary structures of TeCPC G29R and TeAPC G21R were analyzed by size-exclusion chromatography using a column (about 24 mL, Superdex 200 10/300 GL, GE Healthcare) equipped with Nexera lite inert (SHIMAZU). The size exclusion column was equilibrated with Buffer T at a flow rate of 0.50 mL/min. An aliquot of approximately 20 μL containing 5 μM (about 0.5 mg/mL) trimer proteins in 20 mM Tris-HCl buffer (pH 8.0) was injected onto the column, and the elution was monitored using UV/VIS detectors.

### 2.7. Thermal denaturation

The temperature dependence of fluorescence intensity was measured using the StepOne real-time PCR system (Thermo). The sample volume was set to 20 µL, and ROX was used as the reporter. The scan range was from 25°C to 95°C. The melting temperature (Tm) values were evaluated by plotting the derivative of the temperature-dependent fluorescence intensity with respect to temperature.

Differential scanning calorimetry (DSC) measurements were carried out using a MicroCal PEAQ DSC instrument (Malvern Panalytical). All samples and buffers were thoroughly degassed under vacuum prior to measurements. The protein solution was loaded into the sample cell, and the corresponding buffer was used in the reference cell. Thermal scans were performed over a temperature range of 25 to 95 °C at a scan rate of 0.5 °C/min. Buffer–buffer baseline scans were subtracted from sample thermograms. Data were analyzed using MicroCal PEAQ DSC Analysis Software. Tm was determined from the maximum of the excess heat capacity (Cp) curve. In both thermal denaturation experiments, the protein concentration was set to 5 µM.

### 2.8. Mass spectrometry

The protein solutions in the 20 mM Tris-HCl buffer (pH 8.0) were used for mass spectrometry. The solutions were injected into an ultra-performance liquid chromatography system (ACQUITY UPLC H-Class PLUS system, Waters), in tandem with a quadruple time-of-flight (Q-TOF) mass spectrometer (Xevo G2-XS, Waters). Intact proteins were desalted by chromatographic separation on MassPREP Micro Desalting Column (Waters) with the gradient from H_2_O to acetonitrile, containing 0.1% formic acid. An electrospray ionization (ESI) technique was employed in the positive ion mode. Leucine enkephalin and NaI were used for the lock-mass correction and calibration, respectively. Data were analyzed using the software MassLynx and UNIFI version 1.9.

### 2.9. X-Ray crystallography

The crystallization was conducted by the hanging-drop vapor diffusion method at 291 K. The protein solution of 20 mg/mL in 5 μL was mixed with the same volume of reservoir solution containing 0.1 M Tris-HCl buffer (pH 8.0) and 1.2 M ammonium sulfate for TeCPC G29R or 0.1 M 2-(N-morpholino)ethanesulfonic acid (MES) buffer (pH 6.0) and 1.1 M ammonium sulfate for TeAPC G21R. X-ray measurement was carried out at beamline BL5A in the KEK Photon Factory (Tsukuba, Japan). Glycerol was used as cryoprotectant at the final concentration of 33% (w/v). The dataset was collected using a monochromatic X-ray beam (λ = 1.00 Å) at 100 K. The oscillation angle was 1.0° per image. A total of 180 images were collected. X-ray diffraction data were integrated and scaled using the XDS programs [17]. Data collection and refinement statistics of X-ray diffraction studies are summarized in Table S3. Initial phases were obtained by molecular replacement using Phaser [18], with the previously reported structures (PDB IDs: 3L0F and 3DBJ) as search models. The crystallographic structures were refined by the program PHENIX [19]. Structural figures were prepared using PyMOL [20].

## 3. Results

### 3.1. Engineering and characterization of trimeric TeCPC and TeAPC

Structurally defined trimers were generated by introducing glycine-to-arginine substitutions near the hexamer interface in both TeCPC (G29R) and TeAPC (G21R). For TeCPC, structure-based screening was conducted to identify mutations that would support the hexamer-suppressing G29R substitution and promote trimer formation. Screening of 14 primary TeCPC variants using size-exclusion chromatography of crude extracts identified three mutations at position 121 of the α-subunit (T121L, T121M, and T121Y) that produced significant peak shifts relative to the primary construct, with retention times of 21.1, 21.4, and 20.0 minutes, respectively, compared with 22.7 minutes for the primary TeCPC (Fig. S2). Among these, T121Y produced the largest shift, indicating the greatest enhancement of trimer formation, and was therefore combined with the G29R substitution to generate the final TeCPC variant.

The engineered variants TeCPC G29R, which also contains the T121Y substitution in the α-subunit, and TeAPC G21R were expressed in *E. coli* and purified to homogeneity. Their oligomeric states were examined by size-exclusion column chromatography. For both proteins, the major elution peak appeared between the bovine serum albumin (BSA) monomer (67 kDa) and dimer (134 kDa) peaks, consistent with trimer formation (Fig. 1A and B). Serial dilution of the injected protein revealed distinct behaviors for the two complexes. For TeCPC G29R, decreasing concentration resulted in the appearance of an additional peak at an elution volume corresponding to the monomer. In contrast, TeAPC G21R maintained a single elution peak even at concentrations near the detection limit, indicating greater stability of the TeAPC G21R trimer under dilute conditions. This result is consistent with the G21R substitution alone being sufficient to stabilize the trimeric assembly of TeAPC, without requiring an additional trimer-promoting mutation.

**Fig. 1.**
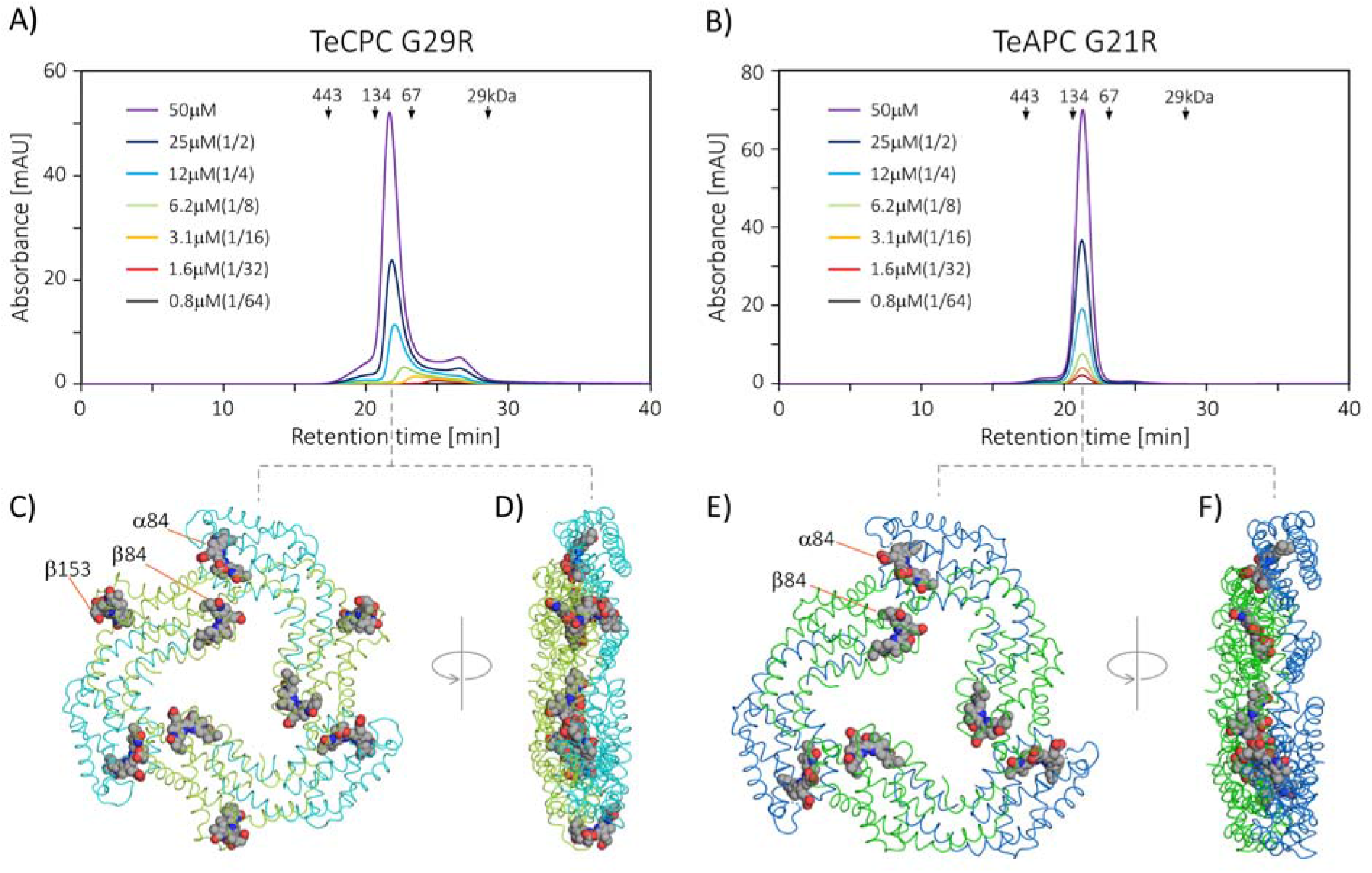
Trimeric structures of TeCPC G29R and TeAPC G21R. (A, B) Size-exclusion column chromatography profiles. 20 μL of serially diluted samples were applied to the column. Sample concentrations are indicated in the legend. Arrows indicate elution times of standards: apoferritin (443 kDa, 17.32 min), dimer of bovine serum albumin (BSA) (134 kDa, 20.60 min), BSA (67 kDa, 23.14 min), and carbonic anhydrase (29 kDa, 28.50 min). (C) Cartoon representation of the loop structure of TeCPC G29R determined by X-ray crystallography. The α- and β-chains are colored cyan and yellow-green, respectively. (D) Side view of TeCPC G29R (90° rotation of panel C). (E) Loop structure of TeAPC G21R. The α- and β-chains are colored marine blue and green, respectively. (F) Side view of TeAPC G21R (90° rotation of panel E). Phycocyanobilin (PCB) pigments at positions α84, β84, and β153 are shown using a van der Waals representation.

### 3.2. X ray crystallographic analysis

To further characterize quaternary structure, X□ray crystallographic analyses were carried out (Table S3). Crystals of both TeCPC G29R and TeAPC G21R belonged to space groups with crystallographic three□fold symmetry, with one monomer present in the asymmetric unit. Crystallographic symmetry analysis revealed that both proteins form uniform trimeric assemblies in the crystal lattice, with TeCPC G29R shown in Fig. 1C and D and TeAPC G21R shown in Fig. 1E and F. Analysis of crystal packing showed no interfaces consistent with hexamer formation in either structure [4,5,21–26], despite the high protein concentrations used for crystallization (Fig. S3). These crystallographic results independently confirmed the trimeric architecture inferred from size-exclusion chromatography.

### 3.3. Verification of post translational modifications

Post□ translational modifications were analyzed by intact mass spectrometry (Fig. 2A and B). For TeCPC G29R, the molecular masses of the α□ and β□ subunits were determined to be 18158.0 Da and 21436.7 Da, respectively. These values are consistent with attachment of one phycocyanobilin (PCB) chromophore to the α subunit and two PCBs plus one methylation on the β□ subunit. For TeAPC G21R, the measured molecular masses of the α□ and β□ subunits were 18093.1 Da and 19822.0 Da, respectively, consistent with one PCB and removal of the N□ terminal methionine on the α□ subunit, and one PCB and one asparagine methylation on the β□ subunit. No significant signals corresponding to chromophore deficient subunits were detected, whereas an additional peak at 19055.2 Da was observed but could not be assigned. These assignments were further supported by omit difference electron-density maps (Fig. 2C–H).

**Fig. 2.**
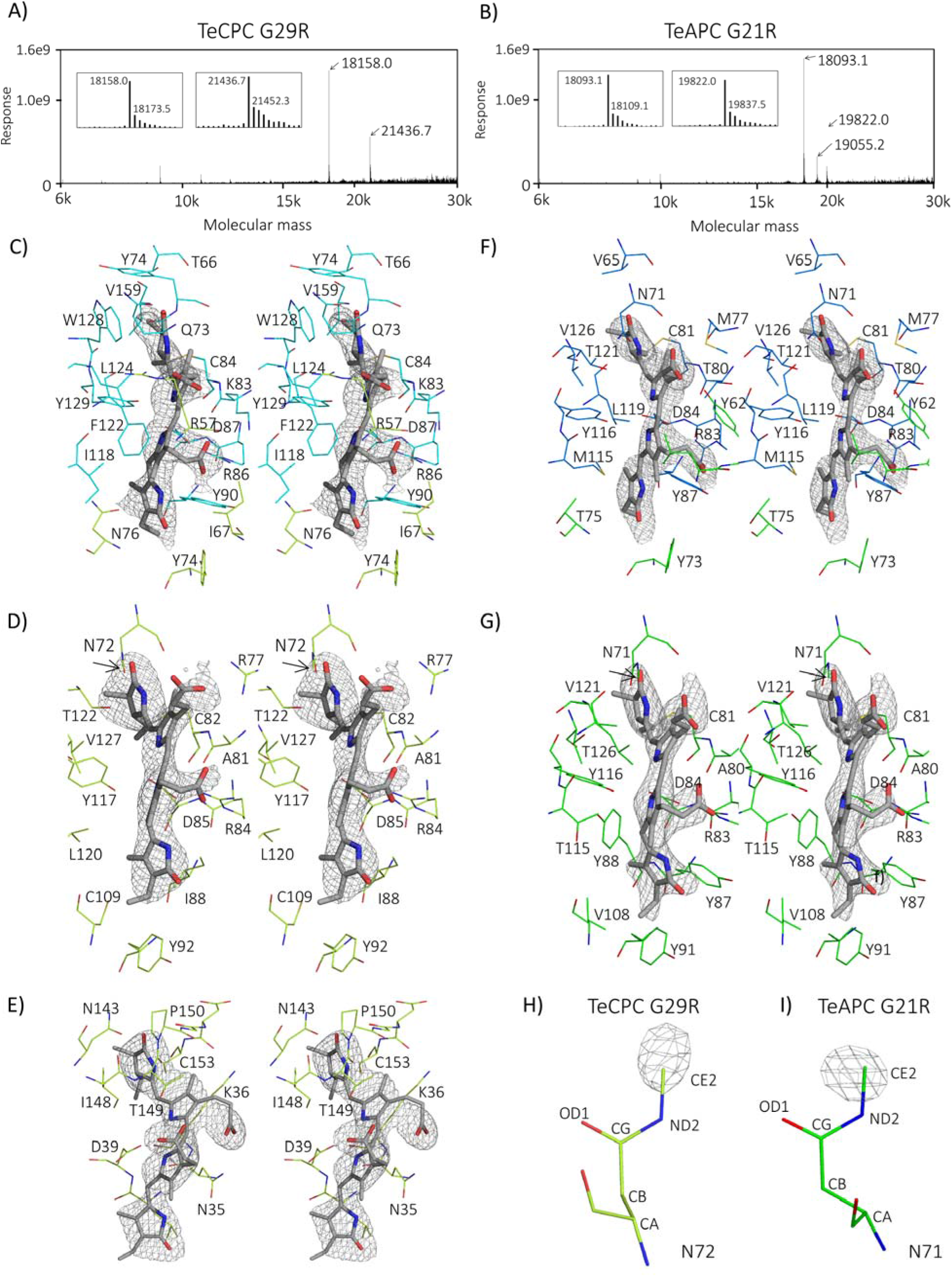
Posttranslational modifications of TeCPC G29R and TeAPC G21R. The intact mass spectrometry analysis for (A) TeCPC G29R and (B) TeAPC G21R. The inset shows the magnified profile around the two significant peaks. (C–G) Difference Fourier electron density maps at 2σ level with wall-eyes stereo view. The maps were calculated after omitting each PCB chromophore individually. PCB models of α84, β84 and β153 for TeCPC G29R are shown in (C), (D) and (E), respectively, and PCB models of α84 and β84 for TeAPC G21R are shown in (F) and (G), respectively. (H, I) Difference electron-density maps around the methylated Asn residue in TeCPC G29R and TeAPC G21R, respectively. The methyl carbon atom was omitted from the model prior to map calculation. The carbon atoms of amino acid residues in the α-chain and β-chain of TeCPC G29R or TeAPC G21R are colored in cyan, yellow-green, marine blue and green, respectively.

### 3.4. Assessment of thermal stability and sample homogeneity

Thermal stabilities of TeCPC G29R and TeAPC G21R were investigated using differential scanning fluorimetry (DSF) and differential scanning calorimetry (DSC) and summarized in Fig. 3. The first-derivative DSF profiles exhibited predominantly single transitions events for both proteins with apparent melting temperatures (Tm) of 69 and 71 °C for TeCPC G29R and TeAPC G21R, respectively. In contrast, DSC revealed more complex thermal behaviors, resolving three distinct transitions for each protein. The baseline-corrected DSC thermograms were therefore analyzed using a three-component deconvolution model, yielding three thermal transitions at 68.5, 71.0, and 77.0□°C for TeCPC G29R, and at 70.0, 75.9, and 83.1□°C for TeAPC G21R, respectively. The presence of multiple calorimetric transitions indicates that the thermal denaturation process cannot be adequately described by a single two-state transition.

**Fig. 3.**
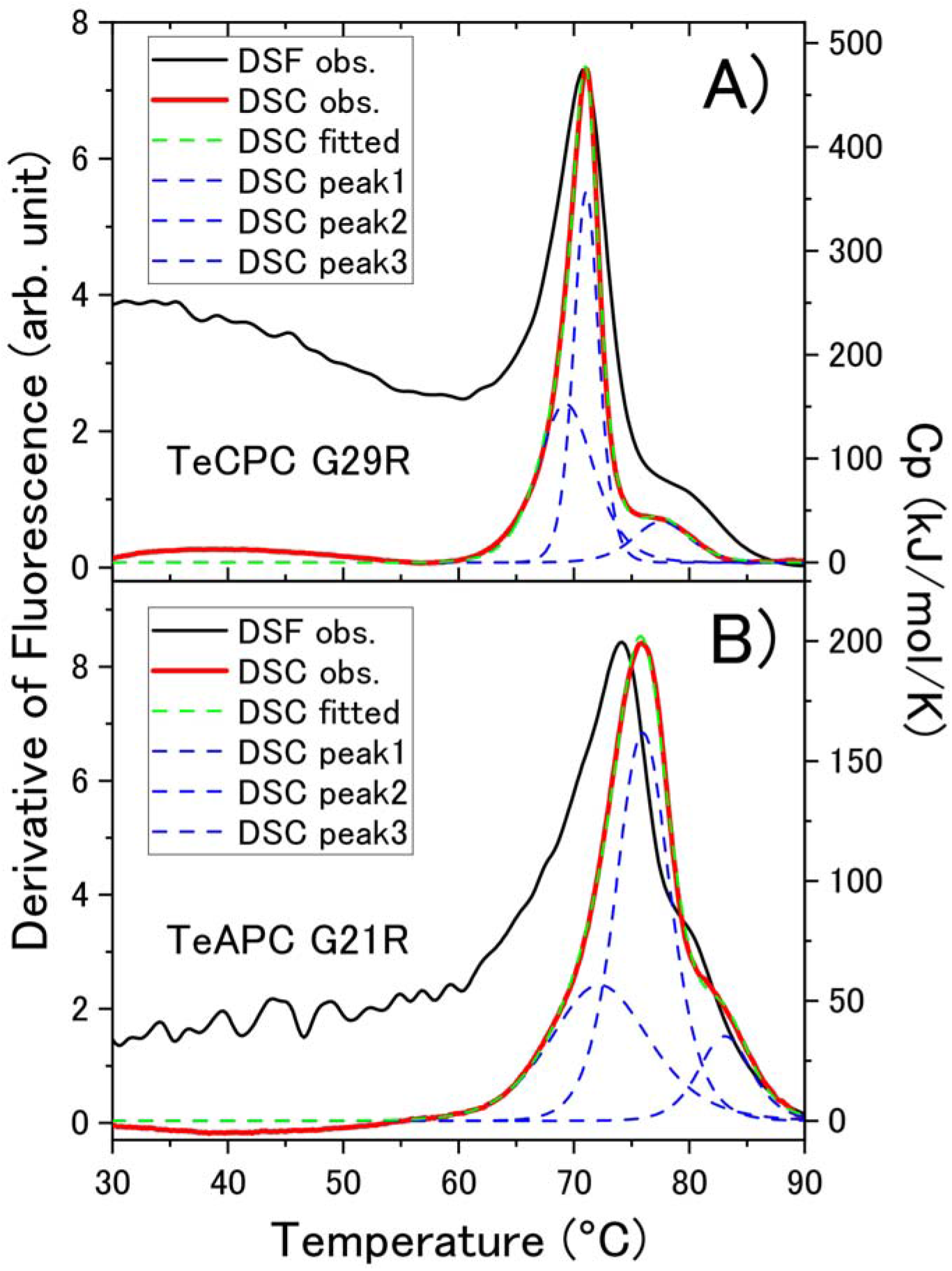
Thermal denaturation profiles of TeCPC G29R and TeAPC G21R. (A) TeCPC G29R and (B) TeAPC G21R. Black solid lines show the first derivatives of the DSF fluorescence signals. Red solid lines represent the baseline-corrected DSC thermograms. The DSC thermograms were analyzed using a three-component deconvolution model. The overall fitted curves and individual transition components are shown as green and blue dashed lines, respectively.

Absorption spectra measured over a range of protein concentrations exhibited no significant changes (Fig. S4), suggesting that oligomeric states were preserved across the conditions tested.

## 4. Discussion

Recombinant protein production in *E. coli* provides a highly efficient approach for introducing amino acid mutations and rational protein design. Many studies have reported co-expression of enzyme systems involved in chromophore biosynthesis and attachment by lyases [27–32]. For physicochemical analyses aimed at understanding the hierarchical organization of efficient light harvesting, it is essential to use samples that are as homogeneous as possible. In this work, the engineering strategy centered on structure-guided substitutions at glycine residues located near the hexamer interface to suppress higher-order assembly. Specifically, we introduced G29R in TeCPC and G21R in TeAPC to create steric hindrance near the C2 symmetry axis that suppresses hexamer formation and stabilizes defined trimeric assemblies. Substitution of these glycine residues (Figs. S5 and S6) with arginine effectively altered oligomerization behavior, as demonstrated by both solution-based and crystallographic analyses. For TeCPC, the additional T121Y substitution in the α-subunit was identified through structure-based screening as a mutation that promotes trimer formation. Unlike TeAPC, G29R alone was insufficient to generate a stable TeCPC trimer. Structural analysis suggests that the T121Y substitution fills a cavity at the inter-subunit interface and strengthens local packing interactions, thereby improving trimer stability. The X-ray crystal structure reveals that the side chain of Tyr121 in the α-subunit interacts with the β-subunit (Fig. S5). Notably, this residue is located away from the hexameric interface, suggesting that its stabilizing effect may arise from local α–β subunit interactions rather than direct suppression of hexameric assembly.

Size□exclusion chromatography showed that both proteins predominantly form trimers, although TeCPC G29R exhibits partial dissociation at low concentration, whereas TeAPC G21R retains a stable trimeric state. This difference suggests intrinsic variations in intersubunit interactions between TeCPC G29R and TeAPC G21R, even in the absence of linker proteins. This observation is also consistent with the finding that TeAPC required only the G21R substitution to achieve stable trimer formation, whereas TeCPC required the additional T121Y mutation. Importantly, the crystallographic analyses provided direct structural evidence for the defined trimeric architecture of the engineered proteins and the absence of detectable hexameric packing interactions in the crystal lattice. The observation that both proteins crystallize with three fold symmetry and lack detectable hexameric contacts, even under high concentration conditions, indicates that the introduced mutations efficiently suppress higher order assembly. The consistency between solution and crystalline states supports the predominance of trimeric assemblies in the engineered complexes.

A key requirement for functional phycobiliproteins is accurate post□translational modification. The combined use of intact mass spectrometry and electron□density analysis of crystallography demonstrated that PCB attachment and Asn methylation were reproduced with high fidelity in the *E. coli* expression system. Importantly, no chromophore□deficient species were detected in intact mass spectrometry, indicating efficient biosynthetic processing. Although an unassigned peak at 19,055.2 Da was observed in the mass spectrum, this signal may arise from a minor co-purified species, and its presence does not affect the assignment of the major subunit peaks or the conclusion of efficient chromophorylation.

Thermal analyses further supported the stability of the engineered complexes (Fig. 3). Similar thermal transitions observed by calorimetry and fluorimetry are consistent with cooperative thermal perturbation of the assembled complexes. In contrast, multiple transitions resolved by differential scanning calorimetry likely reflect thermally distinct structural events. These transitions may arise from dissociation of the trimeric assembly, disruption of αβ heterodimer interactions, and/or unfolding of individual subunits.

Together, these results demonstrate that the engineered TeCPC and TeAPC variants form predominantly trimeric and thermally stable assemblies while faithfully reproducing native post-translational modifications. Importantly, the structural and biochemical validation presented here complements our previous spectroscopic investigations [12], thereby establishing the oligomeric states and post-translational modifications of the engineered complexes with high confidence. These properties establish the complexes as structurally and biochemically defined model systems suitable for future structural, spectroscopic, and mechanistic studies of phycobiliproteins. The engineered trimers provide useful platforms for future investigations of excitation-energy transfer and may serve as building blocks for the development of artificial phycobilisomes and biohybrid light-harvesting systems.

## 5. Conclusion

Structure-guided engineering enabled the generation of structurally defined trimeric forms of TeCPC and TeAPC through suppression of hexamer assembly. Size-exclusion chromatography and X-ray crystallography demonstrated predominantly trimeric architectures, while mass spectrometry confirmed native-like chromophorylation and post-translational modifications. The engineered complexes exhibited high thermal stability and provided structurally and biochemically validated model systems for future structural and spectroscopic investigations of phycobiliproteins. This strategy offers a practical approach for controlling phycobiliprotein quaternary structure and may facilitate the development of artificial phycobilisomes and biohybrid light-harvesting systems.

## CRediT authorship contribution statement

**Motoyasu Adachi**: Conceptualization, Data curation, Investigation, Funding acquisition, Resources, Supervision, Writing – original draft. **Masaaki Tsubouchi**: Funding acquisition, Project administration, Writing – review and editing. **Chie Shibazaki**: Data curation. **Keita Miyake**: Writing – review and editing and **Ryuji Itakura**: Supervision.

## Acknowledgements

We thank Yuji Kagotani and Takemi Kanazawa at QST for their assistance with protein absorption measurements and protein synthesis, respectively. The authors are grateful to Drs. Hidehito Kono and Takatoshi Fujita for valuable discussions and insightful comments.

## Appendix A. Supplementary data

Supplementary data to this article can be found online at ***.

## Data availability

The nucleotide sequences reported in this study have been deposited in the DNA Data Bank of Japan (DDBJ) under accession numbers LC853283, LC853284, LC912691 and LC912692. The atomic coordinates and structure factors have been deposited in the Protein Data Bank (PDB) under accession codes 43EK and 43EL. Other data supporting the findings of this study are available from the corresponding author upon reasonable request.

