## SUPPORTING INFORMATION for "Regulating Light-Harvesting Protein Assembly through Engineered Trimers of Phycocyanin and Allophycocyanin"

### Contents

Fig. S1. DNA and amino acid sequences of primary TeCPC.

Fig. S2. Size-exclusion chromatography analysis of primary TeCPC mutant

Fig. S3. Crystal packings of TeCPC G29R and TeAPC G21R.

Fig. S4. Sample concentration dependence of UV–VIS absorption spectra of TeCPC G29R and TeAPC G21R.

Fig. S5. Interactions between Tyr121 in the α-subunit and the β-subunit.

Fig. S6. Amino acid sequence alignment of CPC and APC.

Table S1 Sequences of oligo DNA primer used for mutation in primary TeCPC

Table S2 List of expression plasmids used in this study.

Table S3 Data collection and refinement statistics (Molecular replacement).

```
GATCTCGATCCCGCGAAATTAATACGACTCACTATAGGGGAATTGTGAGCGGATAACAAT
TCCCCTCTAGAAATAATTTTGTTTAACTTTAAGAAGGAGATATACAT

ATGAAAACCCCGATTACCGAAGCCATTGCCGCCGCCGATACCCAAGGCCGCTTTCTGAGC  60
 M  K  T  P  I  T  E  A  I  A  A  A  D  T  Q  G  R  F  L  S   20

AACACCGAACTGCAGGCGGTGGATGGCCGCTTTAAACGCGCCGTGGCGTCGATGGAGGCG 120
 N  T  E  L  Q  A  V  D  G  R  F  K  R  A  V  A  S  M  E  A   40

GCGCGTGCGCTGACCAACAATGCGCAGAGCCTGATTGACGGTGCCGCGCAGGCGGTGTAC 180
 A  R  A  L  T  N  N  A  Q  S  L  I  D  G  A  A  Q  A  V  Y   60

CAGAAATTCCCGTACACCACCACGATGCAGGGCTCGCAGTATGCCAGCACGCCGGAAGGT 240
 Q  K  F  P  Y  T  T  T  M  Q  G  S  Q  Y  A  S  T  P  E  G   80

AAGGCGAAATGCGCGCGCGATATTGGCTACTACCTGCGCATGGTGACCTATGCCCTGGTG 300
 K  A  K  C  A  R  D  I  G  Y  Y  L  R  M  V  T  Y  A  L  V  100

GCGGGTGGTACCGGCCCTATGGATGAATACCTGATCGCGGGCCTGAGCGAAATCAATAGC 360
 A  G  G  T  G  P  M  D  E  Y  L  I  A  G  L  S  E  I  N  S  120

ACCTTCGACCTGTCGCCGAGCTGGTATATTGAAGCGCTGAAGTACATTAAAGCGAACCAT 420
 T  F  D  L  S  P  S  W  Y  I  E  A  L  K  Y  I  K  A  N  H  140

GGCCTGACCGGTCAAGCCGCGGTGGAGGCCAACGCGTATATCGACTATGCGATCAACGCC 480
 G  L  T  G  Q  A  A  V  E  A  N  A  Y  I  D  Y  A  I  N  A  160

TTATCGTAA
 L  S  *

GCCATACCGCGAAAGGTTTTGCGCCATTCGATGGTGTCCGGGATCTCGACGCTCTCCCTT
ATGCGACTCCTGCATTAGGAAATTAATACGACTCACTATAGGGGAATTGTGAGCGGATAA
CAATTCCCCTGTAGAAATAATTTTGTTTAACTTTAAGAAGGAGATATACC

ATGAAGGATGCGTTCGCCAAAGTGGTGGCCCAAGCCGATGCGCGTGGCGAATTTCTGACC  60
 M  K  D  A  F  A  K  V  V  A  Q  A  D  A  R  G  E  F  L  T   20

AACGCCCAGTTCGATGCGCTGAGCAACCTGGTGAAAGAAGGCAACAAGCGCCTGGATGCC 120
 N  A  Q  F  D  A  L  S  N  L  V  K  E  G  N  K  R  L  D  A   40

GTGAATCGCATTACCAGCAACGCGAGCACCATTGTTGCGAACGCGGCGCGCGCCTTATTT 180
 V  N  R  I  T  S  N  A  S  T  I  V  A  N  A  A  R  A  L  F   60

GCGGAACAGCCGCAACTGATTCAGCCGGGCGGCAACGCCTATACCAACCGCCGTATGGCG 240
 A  E  Q  P  Q  L  I  Q  P  G  G  N  A  Y  T  N  R  R  M  A   80

GCGTGCCTGCGTGACATGGAGATTATTCTGCGCTACGTTACCTACGCGATTCTGGCCGGC 300
 A  C  L  R  D  M  E  I  I  L  R  Y  V  T  Y  A  I  L  A  G  100

GATAGCAGCGTTCTGGATGACCGTTGCCTGTCGGGCCTGCGCGAAACCTACCAGGCCTTA 360
 D  S  S  V  L  D  D  R  C  L  S  G  L  R  E  T  Y  Q  A  L  120

GGTACCCCGGGCAGCTCGGTTGCGGTGGCGATTCAGAAAATGAAGGACGCGGCGATTGCC 420
 G  T  P  G  S  S  V  A  V  A  I  Q  K  M  K  D  A  A  I  A  140

ATCGCGAACGATCCGAGCGGTATCACCCCGGGCGATTGCAGCGCGCTGATGAGCGAAATC 480
 I  A  N  D  P  S  G  I  T  P  G  D  C  S  A  L  M  S  E  I  160

GCGGGCTATTTTGATCGTGCGGCGGCGGCGGTTGCGGGTAAACATCACCATCATCACCAT 540
 A  G  Y  F  D  R  A  A  A  A  V  A  G  K  H  H  H  H  H  H  180

TGGAGCCATCCGCAGTTTGAAAAATGAGATCCGGCTGCTAACAAAGCCCGAAAGG
 W  S  H  P  Q  F  E  K  *
```

Fig. S1. DNA and amino acid sequences of primary TeCPC. Gray letters indicate sequences derived from the pET24a vector. Orange and green boxes represent the T7 promoter and lac operator, respectively. Dark blue and light blue indicate the His tag and StrepII tag, which were added for protein purification. Amino acid residues shown in red indicate the positions targeted for mutational screening to enhance trimer formation.

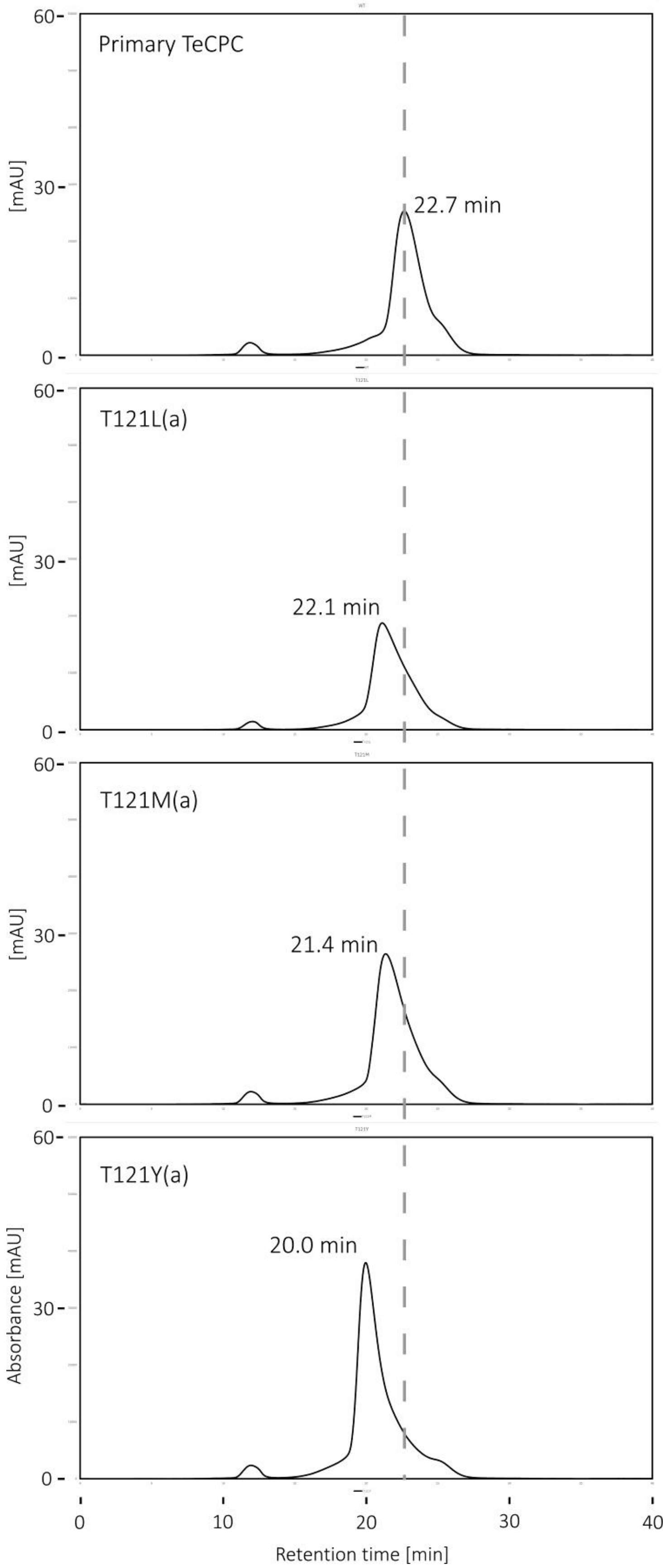

Primary TeCPC
22.7 min
T121L(a)
22.1 min
T121M(a)
21.4 min
T121Y(a)
20.0 min
[mAU]
Absorbance [mAU]
Retention time [min]
0
30
60
0
10
20
30
40

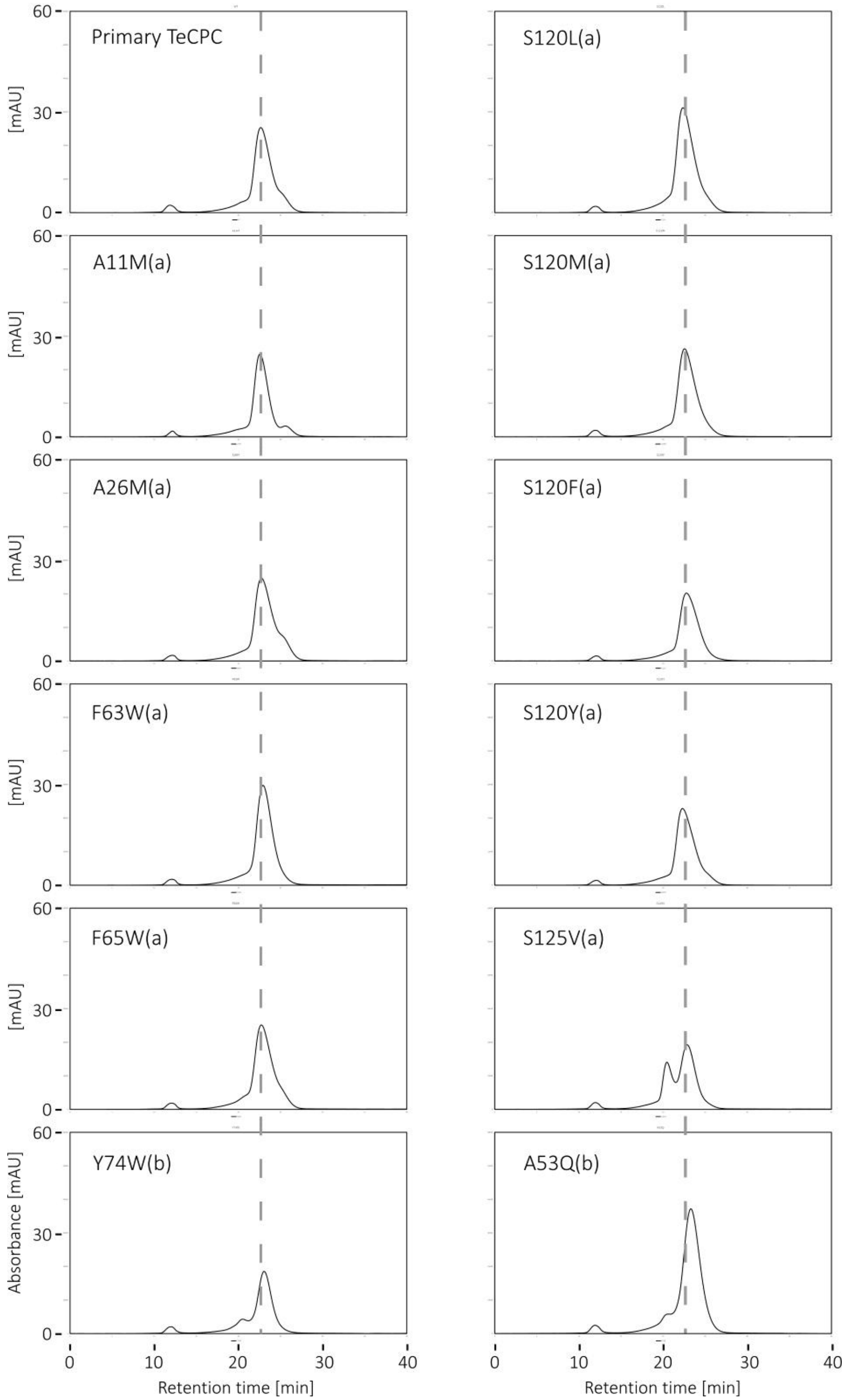

Fig. S2. Size-exclusion chromatography analysis of primary TeCPC mutants. Proteins were expressed in 3-mL cultures. Soluble fractions prepared from sonicated *E. coli* cells were analyzed by size-exclusion chromatography following centrifugation.

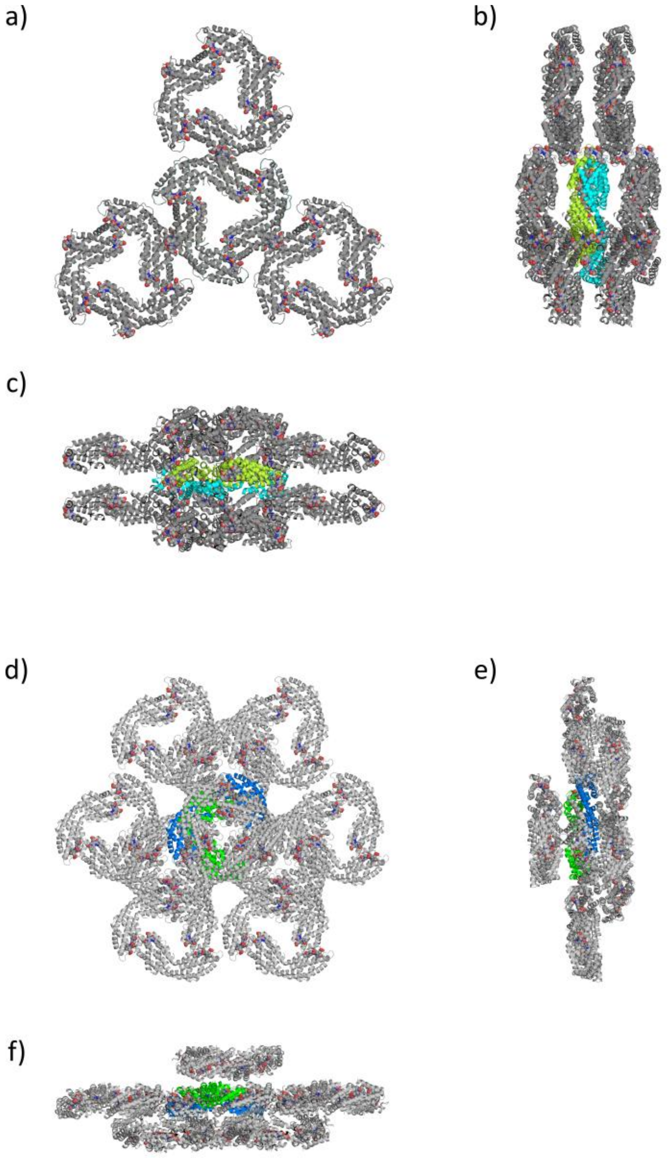

Fig. S3. Crystal packings of TeCPC G29R and TeAPC G21R. Panels a-c show TeCPC G29R, and panels d-f show TeAPC G21R. The α- and β-chains of TeCPC G29R are colored cyan and yellow−green, respectively, whereas the α- and β-chains of TeAPC G21R are colored marine blue and green, respectively. Symmetry-related molecules are shown in gray. Panels b and e show views rotated by 90° about the vertical axis relative to panels a and d, respectively, whereas panels c and f show views rotated by 90° about the horizontal axis. In panel d, the colored molecule is completely obscured by the gray molecule located in front of it. PCB chromophores are shown as ball-and-stick models.

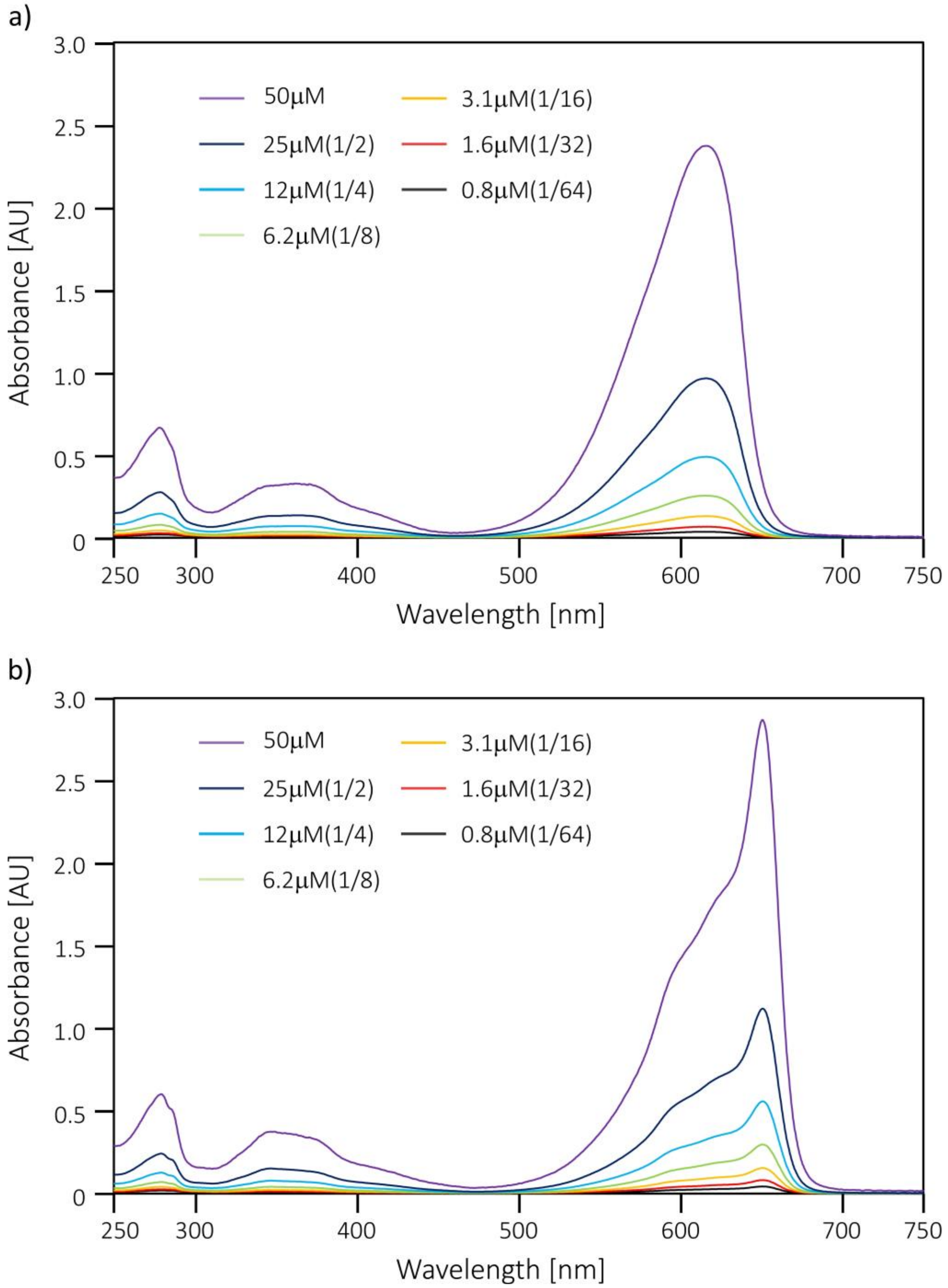

Fig. S4. Sample concentration dependence of UV–VIS absorption spectra of (a) TeCPC G29R and (b) TeAPC G21R. Protein concentrations are indicated in the legend. Prior to size-exclusion chromatography, the protein samples were serially diluted in 20 mM Tris-HCl buffer (pH 8.0) from an initial concentration of 50 μM to 1/64 of the original concentration.

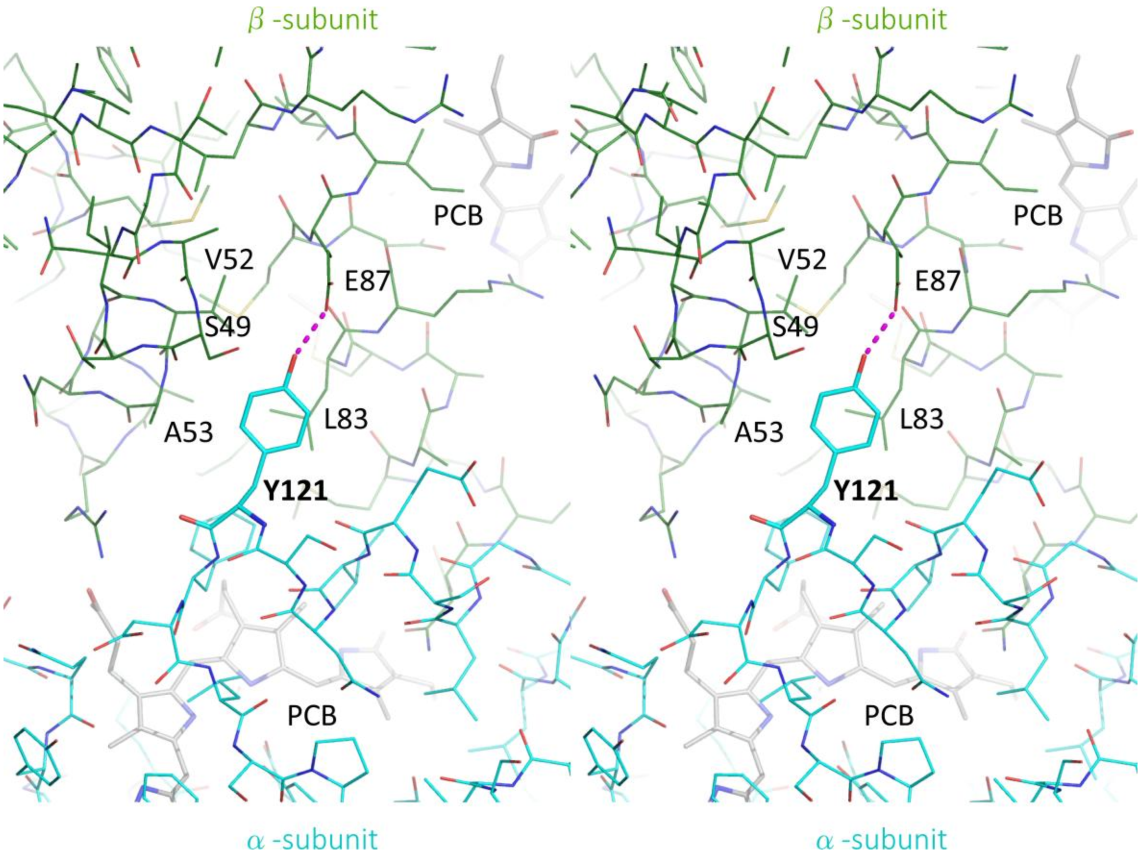

Fig. S5. Interactions between Tyr121 in the α-subunit and the β-subunit (stereo view). The α- and β-subunits are shown as cyan and yellow-green line models, respectively. β-subunit residues Ser49, Val52, Ala53, and Glu87 that interact with Tyr121 are labeled. An intersubunit hydrogen bond is observed between Tyr121 and Glu87, as indicated by a magenta dotted line. The aromatic side chain of Tyr121 is positioned over Leu83.

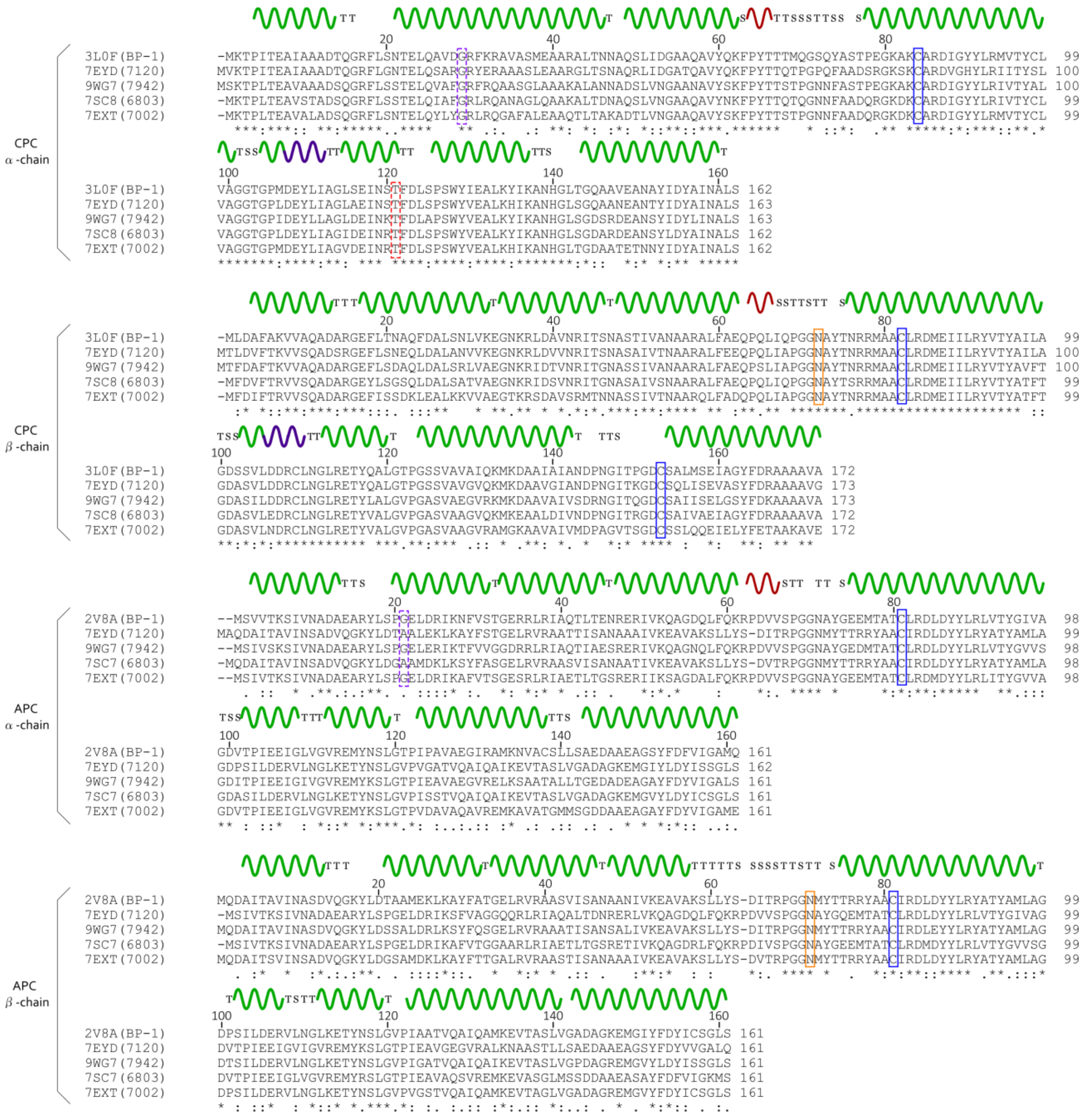

Fig. S6. Amino acid sequence alignment of CPC and APC proteins. Secondary structures were assigned using the DSSP program, with α-helices represented by green lines and $3_{10}$-helices represented by red lines. T and S indicate turns and bends, respectively. Blue boxes indicate chromophore-binding sites, red dashed boxes indicate residues mutated to promote trimer formation, orange boxes indicate methylated asparagine residues, and purple dashed lines indicate residues involved in suppressing hexamer formation.

Table S1. Sequences of oligonucleotide primers used for mutagenesis of primary TeCPC.

| Name | Forward primer* | Name | Reverse primer* |
|---|---|---|---|
| CpcA_A11M_F | ATTGCCATGGCCGATACCCAAGGCCGCTTTCTGAGC | CpcA_A11M_R | ATCGGCCATGGCAATGGCTTCGGTAATCGGGGTTTTC |
| CpcA_A26M_F | CTGCAGATGGTGGATGGCCGCTTTAAACGCGCC | CpcA_A26M_R | ATCCACCATCTGCAGTTCGGTGTTGCTCAGAAAG |
| CpcA_F63W_F | CAGAAATGGCCGTACACCACCACGATGCAGGGC | CpcA_F63W_R | GTACGGCCATTTCTGGTACACCGCCTGCGCGGC |
| CpcA_Y65W_F | TTCCCGTGGACCACCACGATGCAGGGCTCGCAG | CpcA_Y65W_R | GGTGGTCCACGGGAATTTCTGGTACACCGCCTG |
| CpcA_Y74W_F | TCGCAGTGGGCCAGCACGCCGGAAGGTAAGGCG | CpcA_Y74W_R | GCTGGCCCACTGCGAGCCCTGCATCGTGGTGGT |
| CpcA_S120M_F | ATCAATATGACCTTCGACCTGTCGCCGAGCTGG | CpcA_S120M_R | GAAGGTCATATTGATTTCGCTCAGGCCCGCGATC |
| CpcA_S120L_F | ATCAATCTGACCTTCGACCTGTCGCCGAGCTGG | CpcA_S120L_R | GAAGGTCAGATTGATTTCGCTCAGGCCCGCGATC |
| CpcA_S120F_F | ATCAATTTCACCTTCGACCTGTCGCCGAGCTGG | CpcA_S120F_R | GAAGGTGAAATTGATTTCGCTCAGGCCCGCGATC |
| CpcA_S120Y_F | ATCAATTACACCTTCGACCTGTCGCCGAGCTGG | CpcA_S120Y_R | GAAGGTGTAATTGATTTCGCTCAGGCCCGCGATC |
| CpcA_T121L_F | AATAGCCTCTTCGACCTGTCGCCGAGCTGGTAT | CpcA_T121L_R | GTCGAAGAGGCTATTGATTTCGCTCAGGCCCGC |
| CpcA_T121M_F | AATAGCATGTTCGACCTGTCGCCGAGCTGGTAT | CpcA_T121M_R | GTCGAACATGCTATTGATTTCGCTCAGGCCCGC |
| CpcA_T121Y_F | AATAGCTACTTCGACCTGTCGCCGAGCTGGTAT | CpcA_T121Y_R | GTCGAAGTAGCTATTGATTTCGCTCAGGCCCGC |
| CpcA_S125V_F | GACCTGGTGCCGAGCTGGTATATTGAAGCGCTG | CpcA_S125V_R | GCTCGGCACCAGGTCGAAGGTGCTATTGATTTC |
| CpcB_A53Q_F | ATTGTTCAGAACGCGGCGCGCGCCTTATTTGCG | CpcB_A53Q_R | CGCGTTCTGAACAATGGTGCTCGCGTTGCTGGTAATG |

*Underlines indicate overlapping DNA sequences after PCR amplification. Yellow markers denote the codons at the mutation sites.

Table S2. List of expression plasmids used in this study.

| Name of plasmid | Original plasmid | Coded proteins | Organisms* | Accession number (reference sequence) | Accession number deposited |
|---|---|---|---|---|---|
| pET_TeCPC G29R (including T121Y) | pET24a (Novagen) | CpcA | BP-1 | NCBI: WP_011056801 | LC912692 |
| | | CpcB | BP-1 | NCBI: WP_011056800 | |
| pET_TeAPC G21R | pET24a (Novagen) | ApcA | BP-1 | NCBI: WP_011057793 | LC912691 |
| | | ApcB | BP-1 | NCBI: WP_011057792 | |
| pACYC_ HO1PcyA | pACYCDuet-1 (Novagen) | HO1 | 6803 | NCBI: WP_010871494 | LC853284 |
| | | PcyA | 7120 | NCBI: WP_010997850 | |
| pCDF_ MTSEF | pCDFDuet-1 (Novagen) | CpcM | BP-1 | NCBI: WP_011057783 | LC853283 |
| | | CpcT | 7120 | NCBI: WP_010999463 | |
| | | CpcS | 7120 | NCBI: WP_010994793 | |
| | | CpcE | 7120 | NCBI: WP_010994708 | |
| | | CpcF | 7120 | NCBI: WP_010994709 | |

*BP-1: *Thermosynechococcus elongatus* BP-1 (*Thermosynechococcus vestitus* BP-1)

6803: *Synechocystis* sp. PCC 6803

7120: *Nostoc* sp. 7120 (*Anabaena* sp.)

Table S3. Data collection and refinement statistics (Molecular replacement).

| | TeCPC G29R | TeAPC G21R |
|---|---|---|
| **Data collection** | | |
| Space group | $P6_3$ | $P6_322$ |
| Cell dimensions | | |
| *a*, *b*, *c* (Å) | 151.97, 151.97, 39.29 | 101.83, 101.83, 127.17 |
| α, β, γ (°) | 90, 90, 120 | 90, 90, 120 |
| Resolution (Å) | 43.87-2.80(2.95-2.80)* | 47.27-2.54(2.65-2.54)* |
| $R_{merge}$ | 0.205(1.353) | 0.272 (1.907) |
| $I/\sigma I$ | 15.3(2.3) | 17.9(2.2) |
| Completeness (%) | 99.7(99.7) | 100(100) |
| Redundancy | 10.0(10.4) | 18.8(19.4) |
| **Refinement** | | |
| Resolution (Å) | 43.87-2.80 | 47.27-2.54 |
| No. reflections | 13151 | 13422 |
| $R_{work}/ R_{free}$ | 0.202/0.255 | 0.209/0.264 |
| No. atoms | | |
| Protein | 2512** | 2444** |
| Water | 0 | 51 |
| PCB | 129 | 86 |
| B-factors | | |
| Protein | 57.4 | 49.2 |
| Water | - | 47.4 |
| PCB | 56.5 | 45.3 |
| R.m.s deviations | | |
| Bond lengths (Å) | 0.004 | 0.002 |
| Bond angles (º) | 1.518 | 1.254 |
| Ramachandran plot | | |
| favoured region (%) | 94.9 | 92.4 |
| allowed region (%) | 4.5 | 6.6 |
| outlier region (%) | 0.6 | 1.0 |
| PDBID | 43EK | 43EL |

A single crystal was used for structural analysis.

*The highest-resolution shell is shown in parentheses.

**One heterodimer consisting of α- and β-subunits is present in the asymmetric unit.
